# Preclinical Characterization of the Omicron XBB.1.5-Adapted BNT162b2 COVID-19 Vaccine

**DOI:** 10.1101/2023.11.17.567633

**Authors:** Kayvon Modjarrad, Ye Che, Wei Chen, Huixian Wu, Carla I. Cadima, Alexander Muik, Mohan S. Maddur, Kristin R. Tompkins, Lyndsey T. Martinez, Hui Cai, Minah Hong, Sonia Mensah, Brittney Cumbia, Larissa Falcao, Jeanne S. Chang, Kimberly F. Fennell, Kevin Huynh, Thomas J. McLellan, Parag V. Sahasrabudhe, Wei Chen, Michael Cerswell, Miguel A. Garcia, Shilong Li, Rahul Sharma, Weiqiang Li, Kristianne P. Dizon, Stacy Duarte, Frank Gillett, Rachel Smith, Deanne M. Illenberger, Kari E. Sweeney, Annette B. Vogel, Annaliesa S. Anderson, Ugur Sahin, Kena A. Swanson

## Abstract

As SARS-CoV-2 continues to evolve, increasing in its potential for greater transmissibility and immune escape, updated vaccines are needed to boost adaptive immunity to protect against COVID-19 caused by circulating strains. Here, we report features of the monovalent Omicron XBB.1.5-adapted BNT162b2 vaccine, which contains the same mRNA backbone as the original BNT162b2 vaccine, modified by the incorporation of XBB.1.5-specific sequence changes in the encoded prefusion-stabilized SARS-CoV-2 spike protein (S(P2)). Biophysical characterization of Omicron XBB.1.5 S(P2) demonstrated that it maintains a prefusion conformation that adopts a flexible and predominantly open one-RBD-up state, with high affinity binding to the human ACE-2 receptor. When administered as a 4^th^ dose in BNT162b2-experienced mice, the monovalent Omicron XBB.1.5 vaccine elicited substantially higher serum neutralizing titers against pseudotyped viruses of Omicron XBB.1.5, XBB.1.16, XBB.1.16.1, XBB.2.3, EG.5.1 and HV.1 sublineages and the phylogenetically distant BA.2.86 lineage than the bivalent Wild Type + Omicron BA.4/5 vaccine. Similar trends were observed against Omicron XBB sublineage pseudoviruses when the vaccine was administered as a 2-dose primary series in naïve mice. Strong S-specific Th1 CD4^+^ and IFNγ^+^ CD8^+^ T cell responses were also observed. These findings, together with prior experience with variant-adapted vaccine responses in preclinical and clinical studies, suggest that the monovalent Omicron XBB.1.5-adapted BNT162b2 vaccine is anticipated to confer protective immunity against dominant SARS-CoV-2 strains.

**ONE-SENTENCE SUMMARY:** The monovalent Omicron XBB.1.5-adapted BNT162b2 mRNA vaccine encodes a prefusion-stabilized spike immunogen that elicits more potent neutralizing antibody responses against homologous XBB.1.5 and other circulating sublineage pseudoviruses compared to the bivalent Wild Type + Omicron BA.4/5 BNT162b2 vaccine, thus demonstrating the importance of annual strain changes to the COVID-19 vaccine.

## MAIN TEXT

### INTRODUCTION

The evolution of Severe Acute Respiratory Syndrome Coronavirus-2 (SARS-CoV-2), the cause of coronavirus disease 2019 (COVID-19), has been marked by sustained periods of genetic and antigenic drift, best exemplified by the continual emergence of new variants since the appearance of the Omicron variant of concern (VOC) in November 2021 (*1*). The initial antigenic shift to Omicron BA.1, followed by the dominance of Omicron BA.4/5, prompted updates to COVID-19 vaccines to better match prevalent circulating virus strains. Bivalent formulations of the BNT162b2 vaccine, encoding the spike (S) protein of the Wuhan-Hu-1 wild type (WT) strain (GenBank MN908947.3) and either Omicron BA.1 (GISAID EPI_ISL_8880082) or BA.4/5 (GISAID EPI_ISL_15030644) sublineages, subsequently demonstrated effectiveness against COVID-19 in the season after their introduction (*2–5*). The later emergence of recombinant Omicron XBB sublineages, which have dominated the epidemiologic landscape throughout 2023, has since shown that SARS-CoV-2 is able to evolve toward greater transmissibility and to occupy pockets of antigenic space that evade previously established host immunity (*6*). The Omicron XBB.1.5 sublineage exhibits greater antigenic distance from Omicron BA.1 than the latter does from the WT strain (*3, 7*). Waning immunity conferred by prior vaccination or infection with XBB sublineages and the ineffectiveness of nearly all licensed monoclonal antibody therapies against XBB.1.5 (*8*) reflect this immunologic trend (*9, 10*). As such, updating COVID-19 vaccines to more closely matched circulating strains is essential to boosting relevant immunity and maintaining effectiveness against a range of clinical outcomes. Accumulating evidence shows that this principle, well-established for vaccines against influenza and other pathogens, also applies to COVID-19 vaccines (*9, 10*).

BNT162b2 RNA encodes the full-length (FL) S protein stabilized in the prefusion conformation through the substitution of amino acid (aa) positions 986 and 987 to proline residues (S(P2)) (*11–14*), a modification that has increased the antigen’s immunogenicity and expression, as compared to the postfusion state (*15*). To address the increasing dominance of the antigenically distant Omicron XBB sublineages, we modified the original COMIRNATY^®^ vaccine—using the same mRNA backbone as the BNT162b2 that encoded WT S(P2)—to encode an Omicron XBB.1.5 FL S(P2). As the structure of the Omicron XBB.1.5 S(P2) has not been resolved, we sought to characterize the structural and biophysical properties of the mRNA encoded prefusion-stabilized S on this strain-adapted background; including its thermostability profile, human angiotensin converting enzyme-2 (ACE-2) receptor affinity, glycosylation pattern and overall structure and receptor binding domain (RBD) conformational dynamics.

Omicron XBB.1.5-adapted BNT162b2 vaccine formulations were also evaluated in preclinical immunogenicity studies in vaccine-experienced and naïve mice and included the assessment of neutralizing antibody responses against a panel of pseudoviruses of varying phylogenetic proximity and measurement of antigen-specific CD4^+^ and CD8^+^ T cell responses. These studies sought to inform the optimal vaccine valency and composition for eliciting protective immunity. The data presented here provide a basis for understanding the key biophysical and immunologic features of an Omicron XBB.1.5-adapted vaccine in the dynamic landscape of SARS-CoV-2.

## RESULTS

### Omicron XBB.1.5 S(P2) biophysical and structural characterization

The S(P2) antigen of Omicron XBB.1.5 was expressed from DNA corresponding to the XBB.1.5-adapted BNT162b2 RNA coding sequence using similar methods, as previously reported (*13*). After affinity purification, Omicron XBB.1.5 S(P2) eluted as a single peak by size exclusion chromatography (SEC), similar to the WT S(P2) (Fig. 1A). Peak fractions of Omicron XBB.1.5 S(P2) mostly contained cleaved S1 and S2 subunits, as was also observed for WT S(P2) (Fig. S1) (*13*). The SEC peak fraction was then assayed by thermal shift assay (TSA) and biolayer interferometry (BLI). The Omicron XBB.1.5 S(P2) had a melting temperature (T_m_) of 63.0 ± 0.2°C, approximately 4 °C lower than the T_m_ of the WT S(P2) (n = 3) (Fig. 1B). Omicron XBB.1.5 S(P2) bound to the human ACE-2 peptidase domain (ACE-2-PD) with an affinity (K_D_ 4.84 nM) that was slightly less potent than that observed for WT S(P2) (K_D_ 1.24 nM), primarily due to the faster binding off rate of Omicron XBB.1.5 S(P2) (Fig. 1C). The binding affinity of purified monomeric RBD to ACE-2-PD binding affinity was also measured. In this case, Omicron XBB.1.5 RBD exhibited an affinity (K_D_ 1.30 nM) that was 24-fold more potent than that observed for WT RBD (K_D_ 31.3 nM).

**Fig. 1.**
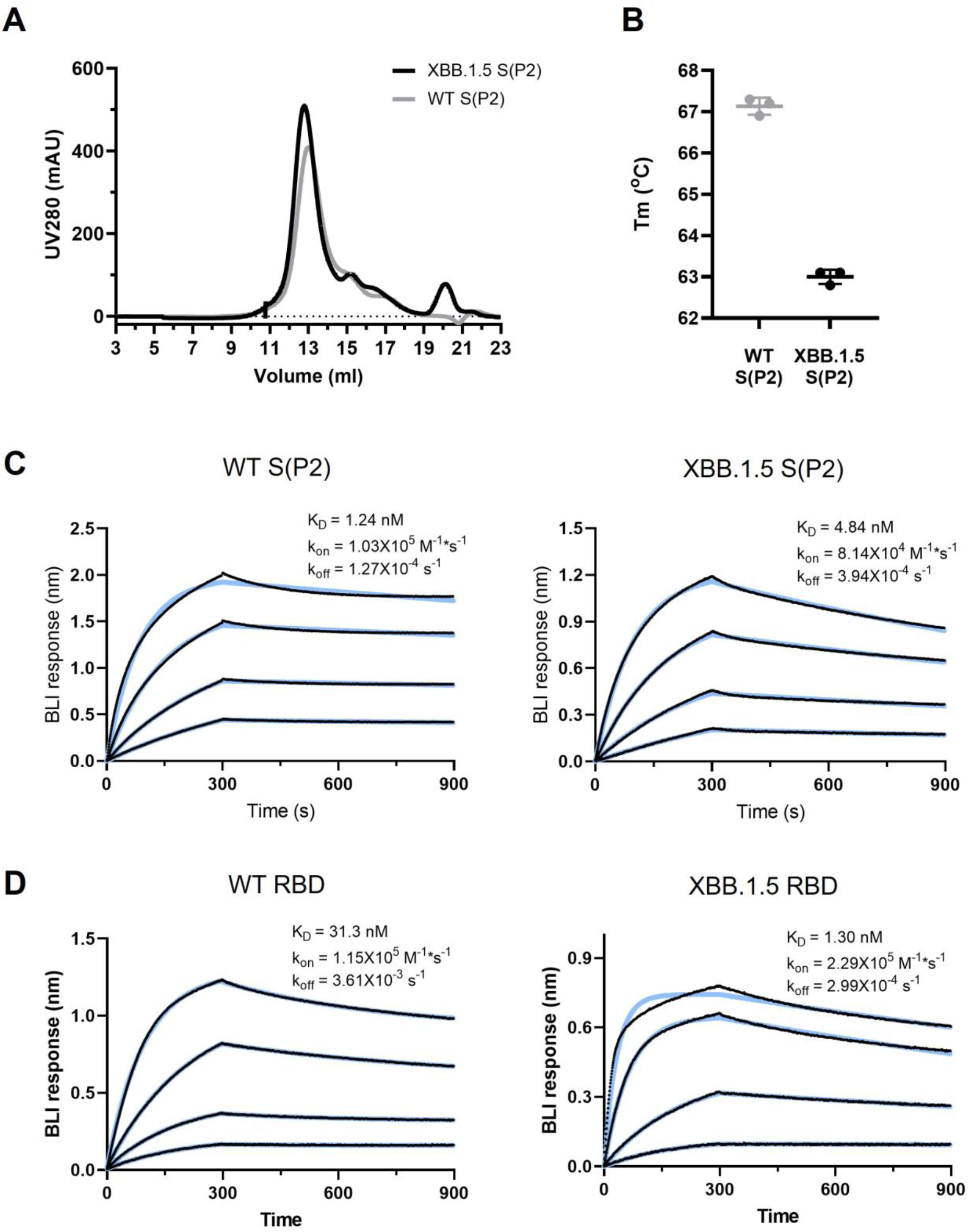
Biophysical Properties and ACE-2 Receptor Binding Affinities SARS-CoV-2 WT and Omicron XBB.1.5 FL S(P2) and RBD. **A.** Size exclusion chromatography (SEC) profile of the purified WT and Omicron XBB.1.5 FL S(P2) proteins equivalent to the protein antigen encoded by the BNT162b2 vaccines. **B.** Melting temperature (Tm) of DDM-purified S(P2) proteins at 0.35 mg/mL concentration. Assay was run in triplicate. **C-D.** Biolayer interferometry (BLI) sensorgram showing binding of (**C**) purified S(P2) proteins and (**D**) RBD to immobilized human angiotensin converting enzyme-2 peptidase domain (ACE-2-PD). Binding data are in black; 1:1 binding model fit to the data is in color. Apparent kinetic parameters are provided in the graph. K_D_ = equilibrium dissociation constant; k_on_ = binding rate constant; k_off_ = dissociation rate constant.

Purified Omicron XBB.1.5 S(P2) was analyzed by liquid chromatography mass spectrometry (LCMS) to identify N-linked glycosylation sites. Twenty-seven N-linked glycosylation sites were detected in S(P2) over a total protein sequence coverage of 92%. The glycosylation pattern was generally similar to that observed for WT S(P2) (*16*) (Fig. S2). However, several new glycosylation sites were also identified in XBB.1.5 S(P2), including N164, N536, N824, N856, N907 and N1119. Two glycosylation sites, N17 and N282, that were reported in WT S(P2) (*16*), were not detected in Omicron XBB1.5 S(P2).

The structure of purified Omicron XBB.1.5 S(P2) was resolved by cryogenic electron microscopy (cryo-EM). Two-dimensional (2D) classification of particles from cryo-EM data revealed a particle population that closely resembled the prefusion conformation of WT S (Fig. 2). Processing and refinement of the dataset (Fig. S3) yielded a high-quality three-dimensional (3D) map with a nominal resolution of 2.98 Å (Fig. 2, Table S1), into which a previously published atomic model (PDB ID: 7TGW) was fitted and rebuilt. The structure revealed that prefusion S(P2) in a 1-RBD-up conformation accounted for the majority (70%) of the high-resolution particles, contrasting with the WT S(P2) all-RBD-down (79.6%) conformation (*13*). The diminished resolution of the RBD-up conformation, as compared to the other parts of the S structure, suggests a conformational flexibility and dynamic equilibrium between RBD ‘up’ and RBD ‘down’ states that is consistent with other reports of SARS-CoV-2 Omicron S structures (*17, 18*). This resolved structure of Omicron XBB.1.5 S(P2), therefore, closely resembles the more open form and flexibility of the S protein of earlier Omicron sublineages (*19*).

**Fig. 2.**
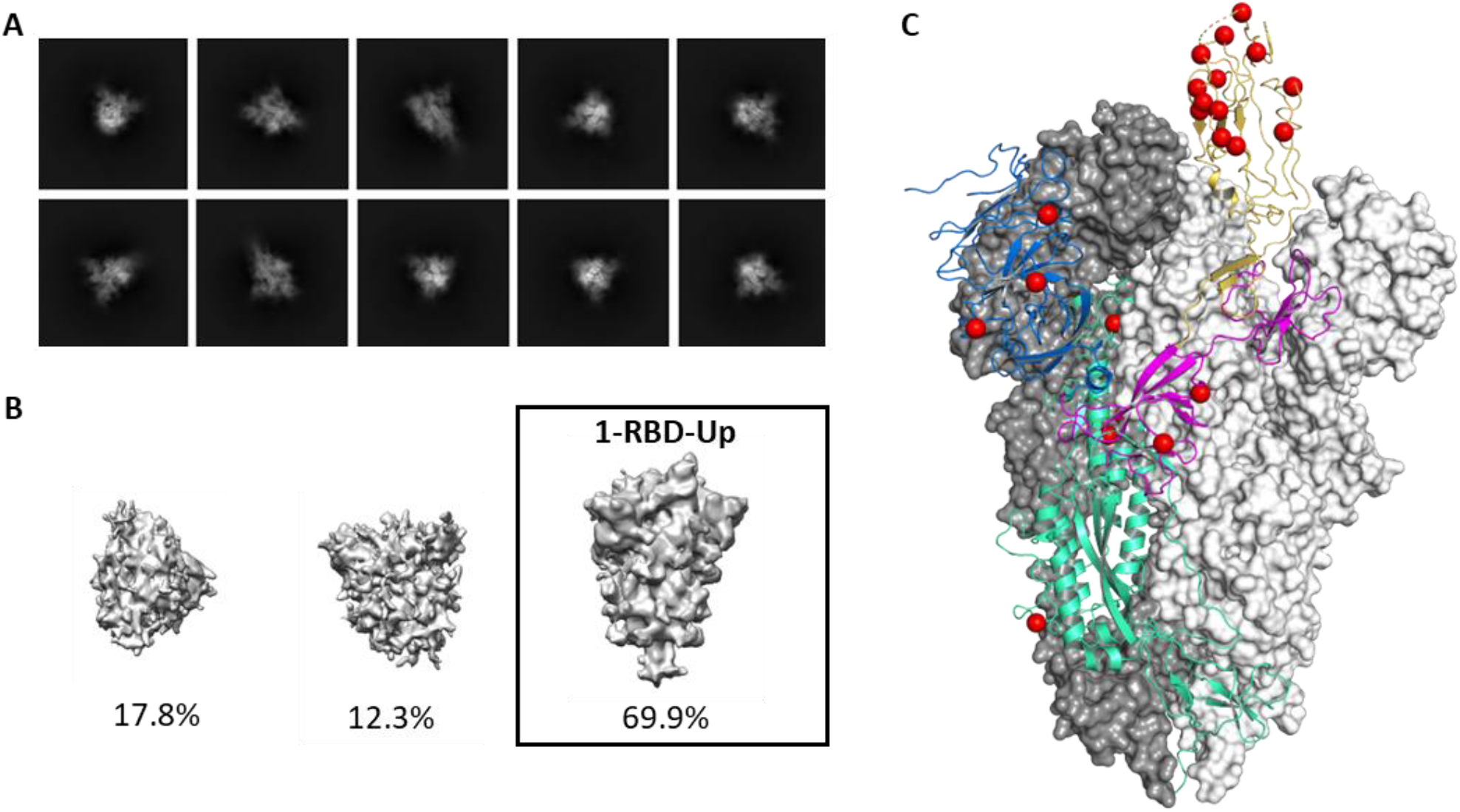
Cryo-EM Structure of SARS-CoV-2 Omicron XBB.1.5 Spike Protein. **A.** Representative 2D class averages of full-length prefusion stabilized Omicron XBB.1.5 S(P2). Box size is 40.5 nm in each dimension. **B.** Maps from *ab initio* reconstruction reveals only one class resembling the S(P2) protein particles with 1-RBD in the ‘up’ position. These particles were used for the final reconstruction. Percentages of the particle population represented in each class are indicated below the models. **C.** The overall structure of Omicron XBB.1.5 S(P2) trimer modeled based on the 2.98 Å density map. Two of the three protomers with RBD in a ‘down’ conformation are represented by a molecular surface colored in white and grey. The remaining protomer with RBD in an ‘up’ conformation is represented by a ribbon diagram; The N-terminal domain is colored blue; the receptor binding domain is colored yellow; the remaining S1 subunit is colored purple; and the S2 subunit is colored green. Amino acid residues that differ between Omicron XBB.1.5 and the ancestral strain are represented by red spheres.

### BNT162b2 Omicron XBB.1.5 immunogenicity

#### Humoral immune response – booster vaccination

Omicron-adapted BNT162b2 formulations were evaluated in two murine studies that varied by prior immune exposure (Fig. S4). In a booster study, female BALB/c mice were experienced with two doses of the monovalent WT BNT162b2 vaccine on Day 0 and Day 21, followed by a single dose of the BNT162b2 bivalent WT + Omicron BA.4/5 vaccine three months later (Fig. S4A). This regimen approximates the relevant immune background of the vaccinated human population that was exposed to S of the ancestral strain and Omicron lineage through vaccination. One month later, animals received one of four BNT162b2 vaccine formulations: monovalent Omicron BA.4/5, bivalent WT + Omicron BA.4/5, monovalent Omicron XBB.1.5 or bivalent Omicron XBB.1.5 + BA.4/5. Sera were collected prior to and one month after the administration of the last dose for assessment of pseudovirus neutralization; splenocytes were collected one month after the last dose to assess T cell responses.

The fifty percent neutralization geometric mean titers (GMTs) at one-month post-4^th^ dose were substantially different across the vaccine groups (Fig. 3A). Neutralizing activity against XBB.1.5 and other circulating XBB sublineages (XBB.1.16, XBB.1.16.1, XBB.2.3, EG.5.1 and HV.1) was highest among animals that received the monovalent Omicron XBB.1.5 booster, particularly compared to the bivalent WT + Omicron BA.4/5 group. GMT values were similar across the Omicron XBB sublineages tested. Neutralizing activity against the genetically divergent BA.2.86 pseudovirus was also highest in the monovalent Omicron XBB.1.5 group, reaching a GMT that was statistically equivalent to the titers for the Omicron XBB pseudoviruses (p > 0.1). Overall, the one-month post-boost GMTs elicited by the monovalent Omicron XBB.1.5 vaccine against the Omicron XBB sublineages were five-to-eight-fold higher than those elicited by the bivalent WT + Omicron BA.4/5 vaccine, while the response against the BA.2.86 lineage was three-fold higher in the monovalent Omicron XBB.1.5 group (Fig. 3B). Two versions of BA.2.86 pseudoviruses were generated and tested because of the variability in spike sequences from early isolates (*20*). These two pseudoviruses, which differed by one amino acid substitution in the subdomain of the S1 subunit (I670V), were equally sensitive to neutralization. As such, data shown for BA.2.86 in Fig. 3 represent the current consensus sequence that does not contain the I670V mutation. Overall, neutralizing antibody responses were highest against WT and Omicron BA.4/5 irrespective of the vaccine group, reflective of prior exposures to WT and Omicron BA.4/5 S.

**Fig. 3.**
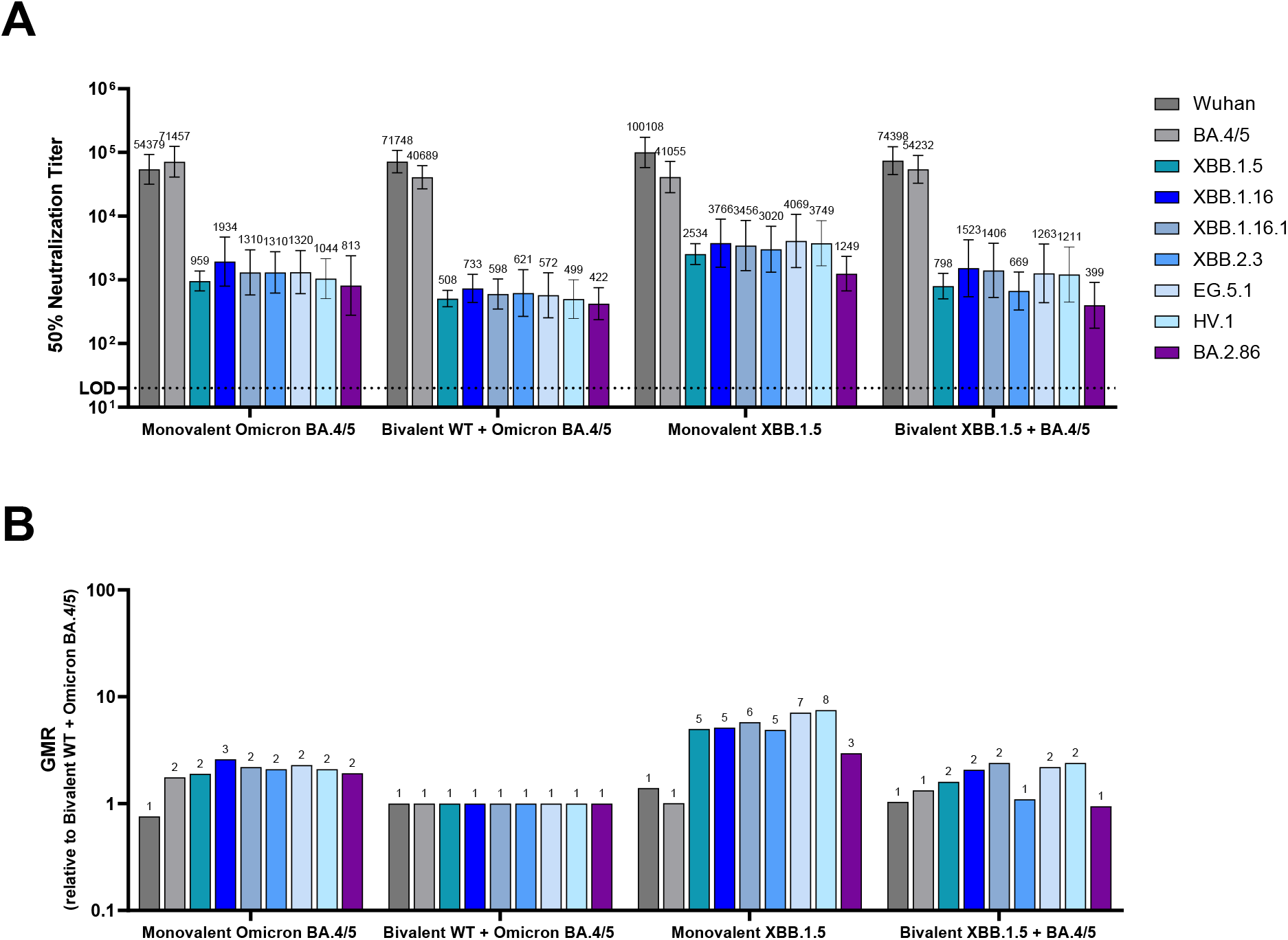
Pseudovirus neutralization titers (NT_50_) elicited by BNT162b2 variant-adapted vaccines administered as a 4_th_ dose in immune-experienced mice. Female BALB/c mice (10/group) that were previously vaccinated (per schedule described in Fig. S4A) with two-doses of monovalent original WT BNT162b2, and one subsequent dose of bivalent WT + Omicron BA.4/5 received a single intramuscular dose of one of these vaccine regimens: monovalent Omicron BA.4/5, bivalent WT+ Omicron BA.4/5, monovalent Omicron XBB.1.5 or bivalent Omicron BA.4/5 + Omicron XBB.1.5. All vaccine formulations contained a total dose of 0.5 µg. Fifty-percent geometric mean serum neutralizing titers were characterized in a pseudovirus neutralization assay at one-month post-4^th^ dose against the WT reference strain, the Omicron sublineages BA.4/5, XBB.1.5, XBB.1.16, XBB.1.16.1, XBB.2.3, EG.5.1, HV.1 and the lineage BA.2.86. A. 50% pseudovirus neutralization titers are shown as geometric mean titers (GMT) ± 95% CI of 10 mice per vaccine group. B. The geometric mean ratio (GMR) is the GMT of individual pseudovirus responses of each vaccine group (monovalent Omicron BA.4/5, monovalent Omicron XBB.1.5 or bivalent Omicron BA.4/5 + Omicron XBB.1.5) divided by the GMT of analogous pseudovirus responses of the bivalent WT + Omicron BA.4/5 group. The limit of detection (LOD) is the lowest serum dilution, 1:20.

The geometric mean titer fold rise (GMFR) in neutralizing activity against Omicron XBB.1.5 pseudovirus from the pre-to post-boost time points were highest in the monovalent XBB.1.5 and bivalent XBB.1.5 + BA.4/5 vaccine groups (GMFR 14.1 and 12.1, respectively) followed by the monovalent Omicron BA.4/5 group (GMFR 10.3) (Fig. S5).

#### Humoral immune response – primary series vaccination

In a primary series study the same vaccine formulations used in the booster study were administered on Days 0 and 21 in naïve female BALB/c mice (Fig. S4B). Sera collected one month after the second dose were tested against the same pseudovirus panel used in the booster study. Similar to the findings of the booster study, the monovalent Omicron XBB.1.5 vaccine elicited substantially higher neutralizing titers against all tested XBB sublineages (XBB.1.5, XBB.1.16, XBB.1.16.1, XBB.2.3 and EG.5.1) (Fig. 4A, 4B) compared to the bivalent WT + Omicron BA.4/5 vaccine, though at much higher titers than those observed in the booster study. Responses were equivalent across the XBB pseudoviruses in the monovalent XBB.1.5 vaccine group (p > 0.1). The BA.2.86 pseudovirus, however, almost completely escaped neutralization in all vaccine groups, except for the bivalent Omicron XBB.1.5 + BA.4/5 where neutralizing responses were an order of magnitude higher than in the other groups.

**Fig. 4.**
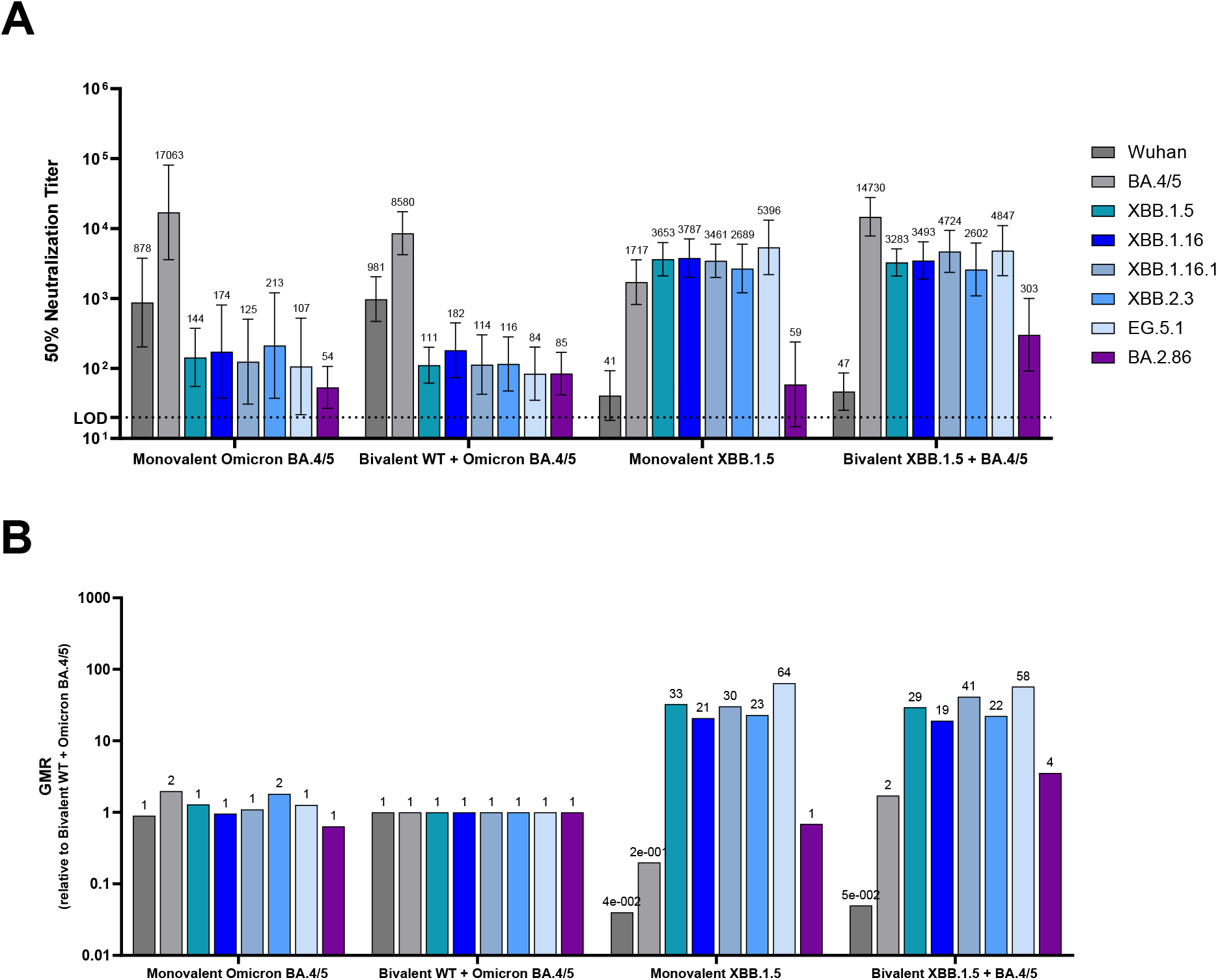
Pseudovirus neutralization titers (NT_50_) elicited by BNT162b2 variant-adapted vaccines administered as a primary series in naïve mice. Female BALB/c mice (10/group) vaccinated with two-doses of one of the following vaccine regimens at a twenty-one-day interval: monovalent Omicron, BA.4/5, bivalent WT + Omicron BA.4/5, monovalent Omicron XBB.1.5 or bivalent Omicron BA.4/5 + Omicron XBB.1.5. All vaccine formulations contained a total dose of 0.5 µg. Serum neutralizing antibody responses were measured by a pseudovirus neutralization assay at one-month post-second dose against the WT reference strain, the Omicron sublineages BA.4/5, XBB.1.5, XBB.1.16, XBB.1.16.1, XBB.2.3, EG.5.1 and the lineage BA.2.86. A. 50% pseudovirus neutralization titers are shown as geometric mean titers (GMT) ± 95% CI of 10 mice per vaccine group. B. The geometric mean ratio (GMR) is the GMT of individual pseudovirus responses of each vaccine group (monovalent Omicron BA.4/5, monovalent Omicron XBB.1.5 or bivalent Omicron BA.4/5 + Omicron XBB.1.5) divided by GMTs of analogous pseudovirus responses of the bivalent WT + Omicron BA.4/5 group. The limit of detection (LOD) is the lowest serum dilution, 1:20.

#### Cellular immune response

T cell responses were measured following XBB.1.5-adapted vaccine administration in both the booster and primary series dosing regimens. Spleens collected one month following the 4^th^ and last vaccine dose were analyzed for frequencies of S-specific T cells, using a flow cytometry-based intracellular cytokine staining (ICS) assay (Fig. S6). Splenocytes were stimulated with S peptide pools representing the amino acid sequence of the WT strain or the Omicron BA.4/5 and Omicron XBB.1.5 sublineages. In the booster study, all vaccine formulations induced high frequencies of S-specific CD4^+^ and CD8^+^ T cells, with a trend toward slightly higher responses in the monovalent Omicron XBB.1.5 group (Fig. 5). The magnitude of IFN-γ-producing T cell responses was higher for CD8^+^ T cells than for CD4^+^ T cells (Fig. 5A, 5B). The frequency of IL-2-and TNF-α-expressing CD4^+^ T cells trended slightly higher than the frequency of CD4^+^ T cells producing IFN-γ (Fig. 5B-D). Very low levels of IL-4-and IL-10-expressing CD4^+^ T cells were observed (Fig. 5E, 5F), thus supporting a Th1-biased response profile that was consistent with prior preclinical and clinical data for BNT162b2 (*13, 21*). T cell responses in the primary series study were similar to those in the booster study, despite the overall magnitude of responses being lower (Fig. 6A-E). Notably, the magnitude of T cell response to each of the strains (WT, Omicron BA.4/5 and Omicron XBB.1.5) was similar within each vaccine group in both booster and primary series studies. These results suggest that polyclonal T cell responses are maintained in mice after primary or booster vaccination and are not significantly impacted by the mutational differences of the Omicron XBB.1.5 sublineage as compared to earlier strains.

**Fig. 5.**
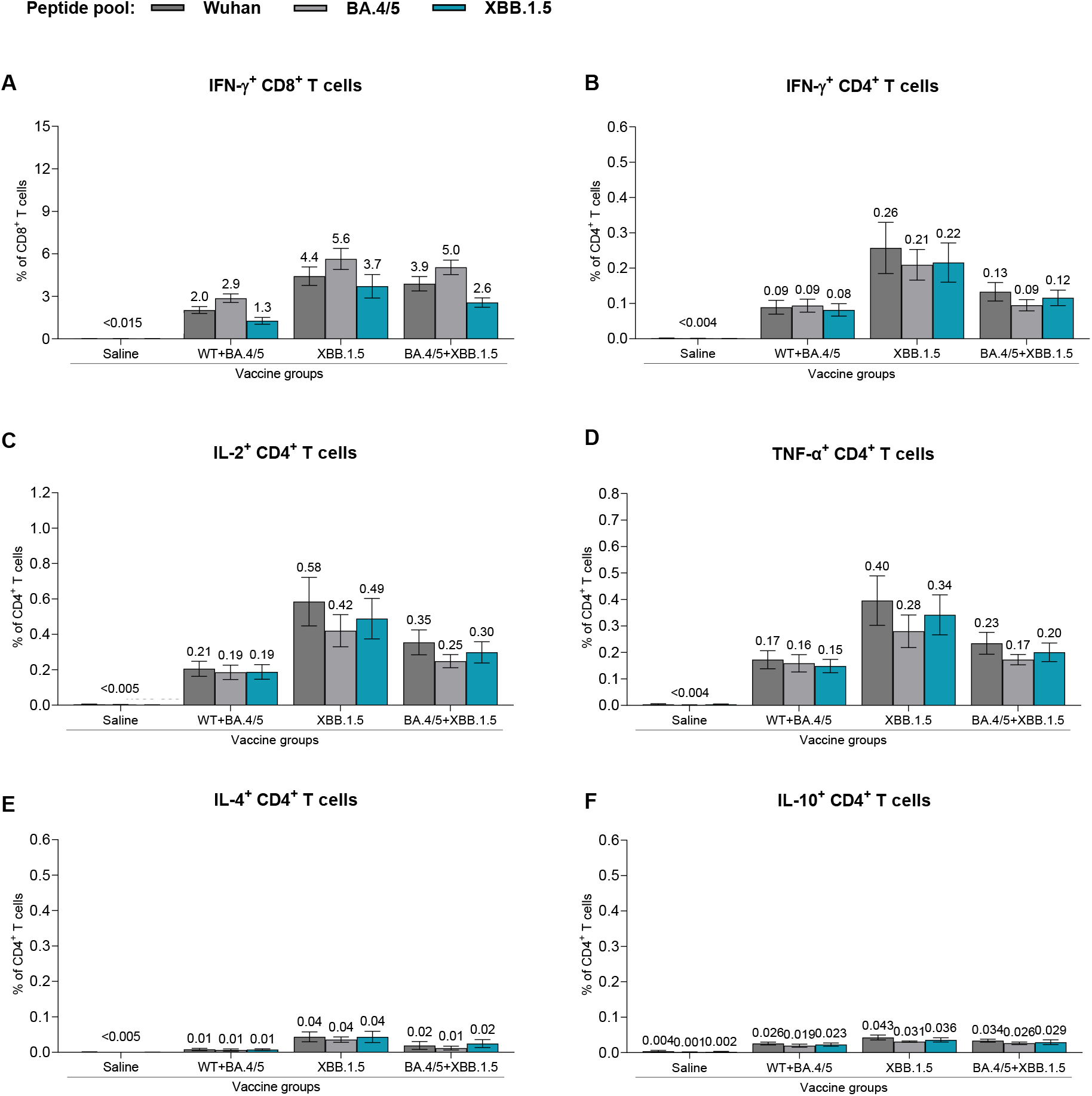
T cell responses elicited by BNT162b2 variant-adapted vaccines administered as a 4_th_ dose in BNT162b2-experienced mice. One-month after the 4th dose, S-specific CD4+ and CD8+ splenocytes (n=5/group) were characterized by a flow cytometry-based intracellular cytokine staining assay. All samples were stimulated separately with S peptide pools from the WT reference strain, Omicron BA.4/5, or XBB.1.5 sublineages. Graphs show the frequency of CD8^+^ T cells expressing IFN-γ (**A**) and the frequency of CD4^+^ T cells expressing IFN-γ (B), IL-2 (C), TNF-α (D), IL-4 (E) and IL-10 (F) in response to stimulation with each peptide pool across vaccine groups. Each symbol represents an individual animal; bars depict mean frequency ± SEM.

**Fig. 6.**
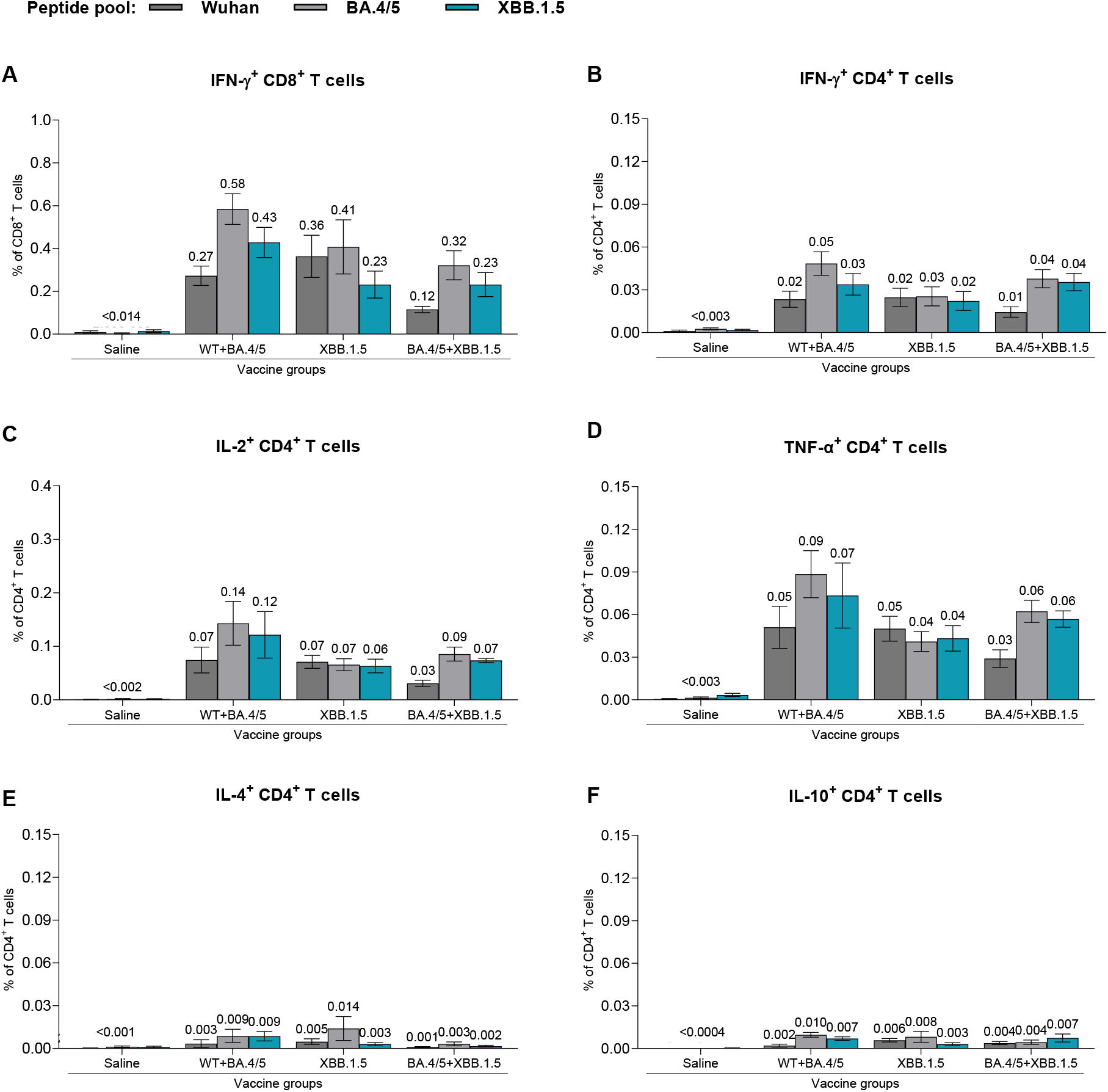
T cell immune responses elicited by BNT162b2 variant-adapted vaccines administered as a primary series in naive mice. At one-month post-second dose (completion of primary series), S-specific T cells from fresh spleens (n=5) were measured by intracellular cytokine staining assay. All samples were stimulated separately with S peptide pools from the WT reference strain and the Omicron BA.4/5 and XBB.1.5 sublineages. Graphs show the frequency of CD8^+^ T cells expressing IFN-γ (**A**) and the frequency of CD4^+^ T cells expressing IFN-γ (B), IL-2 (C), IL-4 (D), TNF-α (E) and IL-10 (F) in response to stimulation with each peptide pool across vaccine groups. Each symbol represents an individual animal; bars depict mean frequency ± SEM.

## DISCUSSION

The evolution of SARS-CoV-2 has prompted an adaptive approach to the continued development of updated vaccines to maintain optimal protection against COVID-19. Since early 2020, when public health crises were declared by multiple national agencies and international normative authorities (*22*), more than 3,400 SARS-CoV-2 unique lineages and sublineages have been identified (*23*). Despite this large genetic diversity, few strains have gained significant advantage to successfully dominate the epidemiologic landscape for extended periods. The most recent strain to exceed a global proportion of 50% is the recombinant Omicron XBB.1.5 sublineage.

Omicron XBB sublineages and their derivatives continue to account for the overwhelming majority of new infections globally (*24*). The descendants of this recombinant lineage cluster consistently exhibit significant immune escape from approved monoclonal antibodies (*8, 25, 26*) The large antigenic distance of these sublineages from earlier SARS-CoV-2 strains, together with waning effectiveness of earlier vaccine iterations based on strains that are no longer circulating and the induction of a more broadly relevant polyclonal antibody response, has necessitated updates to the COVID-19 vaccine.

In the current report, the preclinical data demonstrate an immune response profile that is supportive of the Omicron XBB.1.5 vaccine update. Additionally, for the first time, the trimeric prefusion stabilized structure of Omicron XBB.1.5 S has been resolved. Biophysical characterization studies demonstrate that the Omicron XBB.1.5-adapted BNT162b2 vaccine encodes an S(P2) that authentically presents an antigenically favorable prefusion conformation and ACE-2 binding site. XBB.1.5 S(P2), despite having many mutations, contains biophysical features that are remarkably similar to the S protein of the ancestral WT strain. However, the RBD in Omicron XBB.1.5 S(P2) conforms to a more open and flexible state that contrasts with the closed state of the ancestral strain and early Omicron lineages (*27*). A cryo-EM structure of the BA.2.86 spike was recently reported and showed it adopted a more closed, all-RBD-down conformation that reverts more toward the WT S (*28*). Prior reports have demonstrated that the S of the Omicron BA.2 sublineage, of which XBB.1.5 is a recombinant, may be more compact and thermostable than other variants (*29*). The Omicron XBB.1.5 S(P2), though exhibiting a lower binding affinity for ACE-2-PD than the WT S(P2), has an RBD affinity that is substantially more potent than its ancestral counterpart. The lower T_m_ and less compact structure of XBB.1.5 S(P2) may also result in greater structural instability. These features of the XBB.1.5 S and its components could translate into increased fusion efficiency and account, in part, for the significant growth advantage of this Omicron sublineage. The selective advantage of one conformation versus another, however, remains unclear and raises questions about the optimal positioning of the RBD to best engage the human ACE-2 receptor, while potentially altering the exposure of key regions of the S protein to better escape host immunity.

The observed differences in the N-glycosylation pattern of XBB.1.5 S(P2) compared to the WT S also highlight that the evolution of SARS-CoV-2 is not only driven by amino acid changes and resulting structural conformations but potentially by other post-translational modifications that, for some virus fusion glycoproteins, serve to mask epitopes from antibody recognition. The structural analyses described here may thus inform an understanding of the evolutionary trajectory of SARS-CoV-2, in the context of newer lineages, and in relation to other coronaviruses.

Although a correlate of protection for COVID-19 has not been definitively established, neutralizing antibody titers have trended closely with estimates of vaccine efficacy and effectiveness (*30–32*). Neutralizing antibody responses observed in preclinical animal models have also associated with neutralization trends in clinical studies (*33*). Therefore, assessment of the elicited neutralizing response by the variant adapted vaccines was a paramount objective of the vaccine characterization. When administered as a booster dose or as a primary series in mice, the Omicron XBB.1.5-adapted BNT162b2 vaccine elicited superior neutralizing activity against XBB.1.5 and related XBB sublineage pseudoviruses, including the dominant EG.5.1 and HV.1 strains, compared to that elicited by the bivalent WT + Omicron BA.4/5 vaccine. The data support the conclusion that variant-adapted vaccines offer the ability to maintain optimal immune responses against evolving, circulating SARS-CoV-2 strains.

The more recently emerged Omicron BA.2.86 lineage, a descendant of Omicron BA.2, has approximately sixty and thirty differences in the S amino acid sequence compared to the WT strain and Omicron XBB.1.5 sublineage, respectively. A sequence change of this magnitude has not been observed since the original emergence of Omicron BA.1, which contained approximately thirty amino acid changes relative to Delta, the prior VOC. Despite these sequence changes, the monovalent Omicron XBB.1.5-adapted booster vaccine sera neutralized BA.2.86 to a similar degree as other XBB sublineages, with improved responses over bivalent WT + Omicron BA.4/5 vaccine booster vaccine sera. In contrast, in a naïve background, 2-doses of either the XBB.1.5-adapted or bivalent WT + Omicron BA.4/5-adapted vaccine conferred similarly low neutralizing activity against the BA.2.86 pseudovirus. The bivalent Omicron XBB.1.5 + BA.4/5 vaccine elicited slightly higher BA.2.86 neutralizing titers compared to the other formulations, suggesting a potentially broader coverage of the antigenic space inclusive of where Omicron BA.2.86 resides.

The large discrepancy in BA.2.86 neutralization between the booster and primary series studies indicates that the genetic sequence divergence of this lineage translates into an immunologic difference in a naïve background but does not confer immune escape when the host has multiple prior exposures to antigens that broadly cover the SARS-CoV antigenic space. These data, therefore, demonstrate a significant antigenic distance of this new lineage from preceding ones, though that distance is rendered less important in a population with a diversity of prior immune experience. To date, approximately 1,400 sequences of the BA.2.86 lineage and its derivatives (i.e., BA.2.86.1, JN.1, JQ.1) have been deposited into GISAID since the first confirmed BA.2.86 case (as of July 31, 2023). BA.2.86 remains designated as a variant under monitoring (VUM) by the World Health Organization due to the substantial amino acid changes in its S protein (*34*). However, no sublineage from the BA.2.86 cluster has been reported to cause an increase in COVID-19 disease severity or deaths (*35–41*).

The variant-adapted vaccines evaluated in this study, including the monovalent Omicron XBB.1.5 formulation, elicited robust Th1-type CD4^+^ and IFN-γ-secreting CD8+ T cell responses against S peptide pools representing the FL S of WT, Omicron BA.4/5 and Omicron XBB.1.5.

These findings are consistent with observed trends for multiple variants where antigenic drift and even major shifts to new lineages and sublineages do not substantially erode previously established T cell-mediated immunity (*42, 43*). The likely consequence of a maintained cellular immune response is more durable effectiveness against severe clinical outcomes (*44*).

The findings reported here demonstrate that the monovalent Omicron XBB.1.5-adapted BNT162b2 vaccine encodes a prefusion stabilized S(P2) protein that tightly binds the ACE-2 receptor, maintains a relatively open and flexible conformation, and confers optimal immune responses against contemporaneous SARS-CoV-2 strains. Strengths of this study include the booster immunogenicity study design, which aims to approximate the vaccine-experienced background of the BNT162b2-vaccinated population by pre-exposing animals to both the original monovalent WT vaccine and bivalent WT + Omicron BA.4/5 vaccine. Limitations include the inability to faithfully recapitulate the entire spectrum of immune experience, such as the hybrid immunity gained from prior SARS-CoV-2 infection and vaccination. This immune background likely reflects the majority of the vaccinated population, as seroepidemiology studies show that most individuals have experienced SARS-CoV-2, even among pediatric cohorts (*45*). It was still important to evaluate the Omicron XBB.1.5-adapted vaccine in an immune naïve setting, as there remains a steady proportion of individuals, primarily among the youngest pediatric population, who have not yet been exposed to SARS-CoV-2 through infection or vaccination.

The aggregate data reported here provide a basis for expecting a robust immune response in humans, indicative of a reduction from severe disease outcomes such as hospitalization and death against XBB sublineage infections and resulting COVID-19 disease from ongoing clinical studies (*46*). Preclinical data have reliably predicted responses in humans to vaccination throughout the lifecycle of the original monovalent WT and variant-adapted BNT162b2 vaccines. These types of data now form the basis for regulatory authorizations and approvals of updated formulations, including the bivalent WT + Omicron BA.4/5 vaccine in 2022, and more recently, the monovalent Omicron XBB.1.5 vaccine. COVID-19 epidemiology and immunology continue to be dynamic; as such, safe and effective vaccines will need to keep pace by remaining adaptable to ensure rapid approval and broad access to at-risk populations.

## MATERIALS AND METHODS

### Study Design

The primary aims of the studies reported here are to characterize the biophysical, structural, and immunologic features of the Omicron XBB.1.5 sublineage S(P2) and to test the hypothesis that an adaptation of the BNT162b2 encoded S(P2) to Omicron XBB.1.5 will boost vaccine-elicited immunity to more relevant contemporaneous SARS-CoV-2 strains. The biophysical characterization studies investigated the conformational dynamics of the S RBD and the thermostability profile, affinity to the human ACE-2 receptor, glycosylation patterns and cryo-EM structure of XBB.1.5 FL S(P2). These data were historically compared to the same outputs for the ancestral WT FL S we previously reported (*13*). We evaluated the immunogenicity of Omicron XBB.1.5-adapted BNT162b2 vaccine formulations in both immune-experienced and naïve female BALB/c mice that were vaccinated with monovalent or bivalent SARS-CoV-2 sublineage-modified BNT162b2 vaccines. To evaluate humoral immune responses to the different vaccine series, pseudotyped virus neutralizing serum antibody titers were measured by a pseudovirus neutralization assay (pVNT). To evaluate T cell responses, intracellular cytokine staining (ICS) was performed and quantified by flow cytometry on *ex vivo* S peptide-stimulated splenocytes to detect S-specific, cytokine-secreting T cells.

### Expression and Purification of FL S(P2) and RBD Proteins

In brief, protein sequences of the Omicron XBB.1.5 sublineage and WT (Wuhan-Hu-1) FL S(P2) encoded by BNT162b2 were used to generate a construct containing a C-terminal TwinStrep tag to facilitate affinity purification and were cloned into a pcDNA3.1(+) vector for expression. The isolated RBD constructs contain coding regions from 324-531 (Omicron XBB.1.5) and 327-528 (WT), respectively, of the FL S followed by a C-terminal affinity tag. The FL S(P2) recombinant protein constructs made for this study are summarized in Table S1. Both FL S and RBD protein expressions were conducted in Expi293F cells (ThermoFisher Scientific) grown in Expi293 medium. Upon cell lysis, recombinant FL S(P2) was solubilized in 1% DDM and purified using StrepTactin Sepharose HP resin and size exclusion chromatography (SEC). A modified protocol, based on procedures described previously (*47*), was used for FL S(P2) purification, as detailed in the Supplementary Materials. RBD was expressed as secreted protein and purified via the engineered C-terminal affinity tag.

### Binding Kinetics of Purified FL S(P2) Protein and RBD to Immobilized Human ACE-2-PD

FL S(P2), with a C-terminal TwinStrep tag expressed in Expi293F cells, was detergent solubilized and purified by affinity and size exclusion chromatography. The peak fraction of the purified FL S(P2) and isolated RBD proteins of Omicron XBB.1.5 and WT strains were assessed by biolayer interferometry (BLI) binding to immobilized human ACE-2-PD on an Octet RED384 (FortéBio) at 25 °C in a running buffer that comprised 25 mM Tris pH 7.5, 150 mM NaCl, 1 mM EDTA, and 0.02% DDM, identical to the protein purification buffers. The highest concentration assessed for both FL S(P2) and RBD was 300 nM, with three additional three-fold dilutions. BLI data were collected with Octet Data Acquisition software (version 10.0.0.87) and processed and analyzed using FortéBio Data Analysis software (version 10.0). Binding curves were reference subtracted and fit to a 1:1 Langmuir model to determine binding kinetics and affinity.

### Cryo-EM of Omicron XBB.1.5 FL S(P2)

Purified Omicron XBB.1.5 FL S(P2) was applied to glow discharged Quantifoil R1.2/1.3 200 mesh gold grids and blotted using a Vitrobot Mark IV (ThermoFisher Scientific) before being plunged into liquid ethane cooled by liquid nitrogen. Datasets were collected and analyzed according to details in the Supplementary Materials and as depicted in Fig. S3.

### Additional *In Vitro* Characterization of Omicron XBB.1.5 FL S(P2) Protein

Full details of methods used for the remainder of the *in vitro* characterization of the FL S(P2) protein (*e.g.*, purification and chromatography, mass spectrometry, and thermal shift assay) are provided in the Supplementary Materials.

### Animal Ethics

All murine experiments were performed at Pfizer, Inc. (Pearl River, NY, USA), which is accredited by the Association for Assessment and Accreditation of Laboratory Animal Care (AAALAC). All procedures performed on animals were in accordance with regulations and established guidelines and were reviewed and approved by an Institutional Animal Care and Use Committee or through an ethical review process.

### BNT162b2 mRNA XBB.1.5. Vaccine Modification and Formulation

The XBB.1.5 adapted vaccine encodes the S(P2) of XBB.1.5 (GISAID EPI_ISL_16292655) on the BNT162b2 RNA backbone. Purified nucleoside-modified RNA was formulated into lipid nanoparticles by mixing together an organic phase lipid mixture with an RNA aqueous phase, and subsequently purifying the mix to yield a lipid nanoparticle composition similar to one previously described (*48*).

### Immunogenicity in BNT162b2-Experienced Mice

Female BALB/c mice (10 per group, age 6-8 weeks; Jackson Laboratory) were vaccinated and bled as shown in Fig. S4A. In brief, mice were vaccinated intramuscularly with a 2-dose series (Day 0, 21) of a 0.5 µg dose level of original BNT162b2 WT vaccine, followed by a 3^rd^ dose booster (Day 105) of bivalent WT + Omicron BA.4/5 vaccine, and a 4^th^ dose booster (Day 134) of either monovalent Omicron XBB.1.5, monovalent Omicron BA.4/5, bivalent Omicron XBB.1.5 + BA.4/5 or bivalent WT + Omicron BA.4/5 sublineage-modified vaccines. Bivalent formulations contained equal quantities of each mRNA (0.25 µg each) and a total dose level of 0.5 µg. A control group of ten mice received saline injections according to the same schedule in place of active vaccines. A total volume of 50 µL of vaccine or saline was administered intramuscularly to the upper outer hind leg for each animal. Animals and injection sites were observed immediately after vaccination. Sera were collected for evaluation of pseudovirus neutralizing antibody responses prior to the 4^th^ dose (Day 134) and at the final post-vaccination timepoint (Day 160). Spleens were also collected at Day 160 to evaluate cell-mediated immune responses. When evaluating neutralizing antibody responses for each pseudovirus over time, it was found that pre-boost, baseline GMTs were highest overall against WT, lower against Omicron BA.4/5 and minimal against Omicron XBB.1.5. These trends were similar across vaccine groups (Fig. S7).

### Immunogenicity in Naïve Mice

Female BALB/c mice (10 per group, age 10-12 weeks; Jackson Laboratory) were vaccinated and bled according to the illustration in Fig. S4B. In brief, mice were vaccinated intramuscularly on Days 0 and 21 with either monovalent Omicron XBB.1.5, monovalent Omicron BA.4/5, bivalent Omicron XBB.1.5 + BA.4/5 or bivalent WT + Omicron BA.4/5-adapted vaccines. The control group received saline injections according to the same schedule as active vaccine groups. Sera and spleens were collected 28 days after the second dose (day 49) for evaluation of pseudovirus neutralizing antibody responses and cell-mediated immune responses, respectively.

### Pseudovirus Neutralization Assay

Pseudovirus stocks were generated in HEK-293T cells (ATCC, ref.# CRL-3216) using SARS-CoV-2 spike plasmid DNA and vesicular stomatitis virus (VSV; VSVΔG(G)-GFP virus: Kerafast, ref.# EH1019-PM). Serial dilutions of heat-inactivated murine sera (3-fold) were incubated with pseudovirus (VSVΔG(G)-GFP expressing SARS-CoV-2 S protein) for 1 h at 37 °C before inoculating confluent Vero (ATCC, ref.# CCL81.2) cell monolayers in 96-well plates. Fluorescent virus-infected foci were detected 19-21 h after inoculation with an anti–VSV pAb (Imanis Life Sciences, ref# REA005) and Alexa488-conjugated secondary antibody (Invitrogen, ref# A-11008) and enumerated using a CTL Immunospot Analyzer (Cellular Technology Limited). A 50% neutralization titer (NT_50_) was calculated as the last reciprocal serum dilution at which 50% of the virus is neutralized compared to wells containing virus only. Each serum sample dilution was tested in duplicate. The assay titer range was 20 to 43,740. Any serum sample that yielded a titer >43,470 was prediluted and repeated to extend the upper titer limit; sera that failed to neutralize at the lowest serum dilution (1:20) were reported to have a neutralizing titer of 20 (lower limit of detection, LLOD). VSV-based pseudoviruses used in the assay expressed the S protein from the following SARS-CoV-2 variants: WT (Wuhan-Hu-1, ancestral strain), BA.4/5, XBB.1.5, XBB.1.16, XBB.1.16.1, XBB.2.3, EG.5.1, HV.1 and BA.2.86. Amino acid sequence alignments for all tested pseudoviruses are provided in Fig. S8.

### T-cell Response Assay

Details of the splenocyte isolation and T-cell response assay are provided in the Supplementary Materials. In brief, antigen-specific T cell responses were analyzed from freshly isolated murine splenocytes with a flow cytometry-based intracellular cytokine staining (ICS) assay, comparing unstimulated (DMSO) response to those observed in splenocytes after stimulation with a peptide library. Three individual peptide pools represented the FL S sequences for WT, Omicron BA.4/5, and Omicron XBB.1.5 (JPT, catalog #s PM-SARS2-SMUT10-2, PM-SARS2-SMUT15-1, PM-WCPV-S-1). The ICS assay detected and quantified CD154 (CD40L), IFN-g, TNF-α, IL-2, IL-4, and IL-10, positive CD4^+^ and CD8^+^ T cells. Samples were acquired on a BD LSR Fortessa flow cytometer with BD FACSDiva software and analyzed using BD FlowJo™ software (Version 10.8). Results are shown as a percentage of CD154^+^ and cytokine positive CD4^+^ T cells and CD8^+^ T cells.

## Statistical Analysis

Mouse immunogenicity data were analyzed using GraphPad PRISM software. Statistical comparisons were made on mouse sera GMTs across pseudoviruses within a vaccine group at the last post-vaccination timepoint of each study with a one-way ANOVA on log-transformed data with Dunnett’s multiple comparisons test.

## List of Supplementary Materials

## Materials and Methods

**Fig. S1.** SDS-PAGE of DDM-Solubilized and Purified Prefusion Stabilized Full-length (FL) Ancestral Variant and Omicron XBB.1.5 S(P2) Proteins.

**Fig. S2.** Glycosylation Patterns of Prefusion Stabilized Full-Length WT and XBB.1.5 Spike.

**Fig. S3.** Cryo-EM workflow for construct pSB6534.

**Fig. S4.** Schema for BNT162b2 variant-modified vaccine mouse immunogenicity studies.

**Fig. S5.** Geometric mean fold rise in pseudovirus neutralization titers (NT_50_) from pre-to post-boost with a BNT162b2 variant-adapted vaccine booster immunization. in immune-experienced mice.

**Fig. S6.** Gating strategy for flow cytometry analysis of T cell responses.

**Fig. S7.** Baseline pseudovirus neutralization titers (NT_50_) prior to BNT162b2 variant-adapted vaccine booster immunization in immune-experienced mice.

**Fig. S8.** SARS-CoV-2 Spike amino acid sequence differences across lineages and sublineages for generated pseudoviruses.

**Table S1**. Expression Constructs of FL S proteins.

**Table S2.** Cryo-EM Data Collection, Processing and Refinement Statistics.

References (*47, 49–51*)

## Acknowledgements

We thank Pfizer and BioNTech colleagues for their scientific, programmatic and operational support. We also thank the Pfizer Comparative Medicine department and veterinary staff at Pearl River, NY for their contributions to the *in vivo* studies; the Pfizer Viral Vaccines staff at Pearl River, NY, for their contributions to assay development and implementation, including Marcus Bolton, and for their contributions to virus variant monitoring, including Subrata Saha and Siddartha Mitra; members of Pfizer Discovery Sciences in Groton, CT, specifically Tim Craig, Nicole Nedoma and Honglei Zhao, for their material and technical support on recombinant protein production; and members of the Pfizer Early Bioprocess Development unit in Pearl River, NY, specifically Lei Hu, Tanveer Sandhu, Mindy Wang, Qi Liu, Lynn Phelan and Justin Moran, for their material and technical support. We also thank Robert G. K. Donald for the comprehensive review of and helpful inputs to this manuscript and Christina D’Arco for invaluable writing and editorial support.

## Author contributions

Conceptualization: KM, YC, KAS

Methodology: KM, YC, WC, HW, MSM, KRT, LTM, HC, LF, JSC, KFF, KWH, TJM, PV, WC^†^, MC, MAG, SL, RS, KES, KAS

Data generation and interpretation: KM, YC, WC, HW, CIC, AM, MSM, KRT, LM, MH, SM, BC, LF, JSC, KFF, KH, TJM, PVS, WC^†^, MC, MAG, SL, RS, WL, KPD, SD, FG, RS, DMI, KES, KAS.

Investigation: KM, YC, WC, HW, MSM, KAS

Visualization: YC, WC, HW, MSM, LTM, MH, WL

Supervision: KM, KAS, AVB, ASA, UŞ

Writing – original draft: KM, KAS, YC

Writing – review & editing: KM, YC, WC, HW, CIC, AM, MSM, KRT, LM, MH, SM, BC, LF, JSC, KFF, KH, TJM, PVS, WC^†^, MC, MAG, SL, RS, WL, KPD, SD, FG, RS, DMI, KES, AVB, ASA, UŞ, KAS.

## Data and materials availability

The final full-length XBB.1.5 S(P2) cryo-EM density map and model are deposited in the Electron Microscopy Data Bank (EMDB) and Protein Data Bank (PDB) under accession codes EMD-42524 and PDB ID 8USZ, respectively.

