## Supplemental Materials for "Preclinical Characterization of the Omicron XBB.1.5-Adapted BNT162b2 COVID-19 Vaccine"

[1]Pfizer Vaccine Research and Development, Pearl River, NY, USA.

[2]Pfizer Discovery Sciences; Groton, CT, USA.

[3]BioNTech; Mainz, Germany.

†Viral Vaccines, Pfizer Vaccine Research and Development; Pearl River, NY, USA.

‡Early Bioprocess Development, Pfizer Vaccine Research and Development; Pearl River, NY, USA.

**The file includes:**

Materials and Methods

Figures S1-S8

Tables S1-S2

References *(1-4)*

### Materials and Methods

#### Expression and Purification of FL S(P2)

Expression of proteins was carried out in Expi293F cells (Thermo Fisher Scientific) grown in Expi293 medium. Cells were transiently transfected with S or RBD protein expression constructs in the pcDNA3.1(+) vector. Expression was conducted at 37 °C for 24 hours before adding Expifectamine enhancers (Thermo Fisher Scientific). After addition of enhancers, the temperature was dropped to 32° C and expression was allowed for another 48-72 hours before collecting. A modified protocol of procedures described by Zhang et al (*1*) was used for purification of the SARS-CoV-2 FL S(P2). Briefly, the transfected cells were lysed in a solution containing Buffer A (100 mM HEPES pH 8.0, 150 mM NaCl, 1 mM EDTA), 1% (w/v) *n*-dodecyl-*β*-D-maltopyranoside (DDM, Anatrace), EDTA-free complete protease inhibitor cocktail (Roche), and Pierce Universal Nuclease (Thermo Fisher) at 4 °C for 1 h. After a clarifying spin at 40,000 × g for 45 min, the supernatant was filtered with 0.2 μm filter (Nalgene 78018-24, 1 LL) before batch bound onto StrepTactin HP resin (Cytiva) equilibrated with the lysis buffer at 4 °C for 1 h. Resin was collected by centrifugation at 1000 × g and loaded onto EconoColumn (Bio-Rad) for gravity flow purification. The column was washed with Buffer A containing 0.5% DDM, 10 mM ATP, and 10 mM $MgCl_2$, followed by additional washes with Buffer A and gradually reduced concentrations of DDM (0.5% - 0.02%). FL S(P2) was eluted with Buffer A containing 0.02% DDM and 5 mM d-Desthiobiotin. The protein was further purified by size exclusion chromatography (SEC) on a Superose 6 10/300 column (Cytiva) in a buffer containing 25 mM Tris pH 7.5, 150 mM NaCl, 1 mM EDTA, and 0.02% DDM. DDM-purified FL S(P2) was eluted as a single peak over SEC.

FL S(P2) protein from the SEC peak fractions were analyzed by denaturing PAGE using a 4–15% Criterion TGX Stain-Free Gel (Bio-Rad, Fig. S1), and used in thermostability ($T_m$), biolayer interferometry (BLI), mass spectrometry and cryogenic electron microscopy (cryo-EM) experiments.

**Expression and Purification of RBD**

The RBDs were expressed using Expi293F cells (Thermo Fisher Scientific) grown in Expi293 medium transiently transfected with the RBD expression constructs in pcDNA3.1(+) vector. The RBD constructs contain an N-terminal S protein leader peptide and coding regions from 324-531 (Omicron XBB.1.5) and 327-528 (ancestral strain), respectively, followed by a C-terminal affinity tag as indicated in Table S1. Expression was conducted at 37 °C for 120 hours before the proteins were collected from cell culture medium. The affinity tagged RBDs were purified on affinity purification columns first, subsequently on Superdex200 gel filtration column (Cytiva), and stored in a buffer containing 100 mM Tris pH7.5, 150 mM NaCl, and 10% glycerol.

**Mass Spectrometry Characterization of N-linked Glycosylation**

Mapping of N-linked glycosylation sites was conducted on recombinant purified Omicron XBB.1.5 S(P2) following precipitation using ice cold acetone and incubated overnight at -20 °C. The protein was pelleted, dissolved in 8M urea, and reduced and alkylated prior to proteolytic digestion. Protein samples were digested in three batches using either trypsin, trypsin/Glu-C, or chymotrypsin. Digested peptide pools from all three reactions were subjected to mass spectrometry analysis to achieve a desired sequence coverage (~92%). Peptides were separated from remaining enzymes using a Microcon-10kDa centrifugal filter (MRCPRT010). The supernatant was collected and lyophilized to dryness. For N-linked analysis, digested samples were reconstituted in O18 water. PNGase F and O-glycosidase were added to remove N-linked and O-linked glycosylation.

The treated digests were analyzed on a Thermo QExactive Oribitrap Mass Spectrometer with an EZ-NanoSpray Source and an EZ-nLC 1200. The peptides were chromatographically separated prior to in-line mass spectrometry analysis with a flow rate of 2 mL/min. The samples were also analyzed on a Thermo Fusion Tribrid Mass Spectrometer outfitted with an EZ-nLC 1200 to perform peptide separations. The system was operated in direct injection mode and the peptides were chromatographically separated prior to in-line mass spectrometry analysis with a flow rate of 450 nL/min. Matrix Science MASCOT and Thermo Freestyle were used for data analysis. The N-linked data was searched with the following variable modifications: Deamidated (NQ), Deamidated:18O(1) (NQ), HexNAc (N), Oxidation (M).

**Cryo-EM Analysis of Omicron XBB.1.5 FL S(P2)**

For high resolution structural determination and analysis, FL S(P2) protein of Omicron XBB.1.5, which is equivalent to the antigen encoded by the Omicron XBB.1.5-adapted BNT162b2 vaccine, was subjected to cryo-EM studies. The purified sample in DDM at 5.0 mg/mL were applied onto the glow discharged Quantifoil R1.2/1.3 200 mesh gold grids and blotted using a Vitrobot Mark IV (ThermoFisher Scientific). A data set of 6,690 movies was recorded using EPU from a Titan Krios G2 transmission electron microscope operating at 300 keV equipped with a Falcon 4i direct electron detector and Selectris Energy Filter (ThermoFisher Scientific). Each movie was collected in counting mode with a pixel size of 0.75 Å/pixel, 10 eV slit and a defocus range of -0.6 µm to -2.6 µm for a total dose of 40.0 $e^-/Å^2$. Each data set was imported and processed in CryoSPARC v4.2.1. All movies were adjusted with patch motion correction and patch CTF estimation. Templates were generated from 2D class averages after automative particle picking by blob picker. These 2D class average templates were used for template-based autopicking to pick particles for the rest of the data processing.

From Template Picker, 1,862,402 particles were autopicked and extracted with a box size of 540 pixels. Iterative 2D classification were carried out to select particles with high resolution views of the spike protein (3131,229 particles). Three initial models were generated using all 131,229 particles in *ab initio* reconstruction resulting in only one map containing 91,663 particles that resemble a spike protein with 1-RBD-up. Heterogeneous refinement of the selected particles resulted in only 1-RBD-up structures. Therefore, all 91,663 particles were subjected to homogeneous refinement, followed by non-uniform refinements, which gave the final 1-RBD-up structure with an overall resolution of 2.98 Å. All refinement steps were done with C1 symmetry. The final resolution was calculated from the Fourier Shell Correlation (FSC) curve from the resolution at the 0.143 FSC cutoff.

A model of the Omicron spike protein structure (PDB: 7TGW) (*2*) was fitted into the final cryo-EM structure and was used as a guide for modeling. The atomic model was built in COOT (*3*) and refined using Phenix real space refinement (*4*). The EM density for the RBD in the up position was weakly resolved. Therefore, the RBD was docked into the EM density and rigid body fitted without side chains unless there were clear side chain densities. The final model including the RBDs were refined in Phenix (version 1.20-4459-0000).

**Stability of WT and Omicron XBB.1.5 FL S(P2) by Thermal Shift Assay (TSA)**

Stability of FL S(P2) proteins was measured by Tycho NT.6 (NanoTemper, firmware version: 1.10.3) and the data were analyzed by the Tycho NT.6 software (version: 1.3.2.880). In brief, a 10 μL solution containing 0.35 mg/mL of protein was loaded into a capillary tube and the ratio of tryptophan fluorescence at emission wavelengths of 350 nm over 330 nm and was measured while ramping the temperature from 35 °C to 95 °C using the pre-programmed protocol of the instrument. The inflection temperature for each thermal melting curve was reported by the Tycho NT.6 software.

**Animal blood collection and splenocyte isolation**

For interim blood draws, mice were bled via the submandibular route. Approximately 150 μL of whole blood was collected dropwise directly into microtainer tubes containing serum separators. For terminal blood draws, the entire available blood volume was collected via cardiac puncture. At all blood collection time points, blood tubes remained at room temperature (RT) for at least 30 min prior to centrifuging at 12,300 × g for 3 min. Each serum sample was aliquoted and heat inactivated at 56 °C for 30 min. Samples were stored at -80 °C after testing.

For flow cytometry of murine splenocytes, spleens were collected from five mice per group at the final time point for each study. Spleens were placed in a 100 µm cell strainer (BD Falcon) immersed in 7 mL of complete RPMI (cRPMI: 10% FBS/RPMI; Pen-Strep; Sodium pyruvate) per mouse per well of a 6-well plate. The plates were maintained on ice during transit and before processing for single cell suspension. Spleens were homogenized, subjected to RBC lysis, and passaged through a cell strainer to remove RBCs and clumps.

**Pseudovirus neutralization test (pVNT)**

Details of the pseudovirus neutralization test (pVNT) are provided in the main manuscript. Amino acid sequence alignments for each pseudovirus S are provided in Fig. S8.

**Intracellular Cytokine Staining (ICS) assay for T-cell response evaluation**

Freshly-isolated splenocytes ($2x10^6$ cells/well) were cultured in cRPMI with media containing DMSO only (unstimulated) or specific amino acid (aa) peptide libraries (15aa, 11aa overlap, 1 to 2 µg/mL/peptide) representing the S amino acid sequences of the original SARS-CoV-2 Wuhan strain (WT), Omicron BA.4/5, and Omicron XBB.1.5 sublineages, separately, for 5 h at 37 °C in the presence of protein transport inhibitors, GolgiPlug and GolgiStop. Following stimulation, splenocytes were incubated with fluorescent-conjugated antibodies to the surface proteins CD19, CD3, CD4 and CD8 (25 ± 5 min at RT) followed by fixation and permeabilization and staining for CD154 (CD40L), IFN-γ, TNF-α, IL-2, IL-4, and IL-10 (25 ± 5 min at RT). Ebioscience fixable viability dye eFluor 506 was used exclude dead cells. After staining, the cells were washed and resuspended in flow cytometry (FC) buffer (2% FBS in PBS). Samples were acquired on a BD LSR Fortessa flow cytometer with BD FACSDiva software. Acquired data files were analyzed using BD FlowJo™ software. Results are background (media-DMSO) subtracted and shown as percentage of $CD154^+$ cytokine-expressing $CD4^+$ T cells and $CD8^+$ T cells. The T-cell gating strategy is shown in Fig. S6.

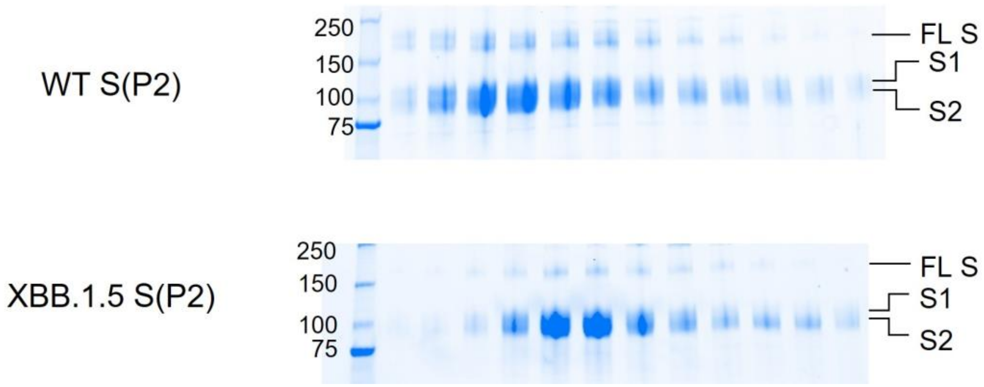


**Fig. S1. SDS-PAGE of DDM-Solubilized and Purified S(P2) WT and Omicron XBB.1.5 S(P2) Proteins.** SDS-PAGE of the SEC fractions from 11.5 mL to 15 mL of the WT S(P2) (upper panel) and 11 mL to 14.5 mL of FL Omicron XBB.1.5 S(P2) (lower panel).

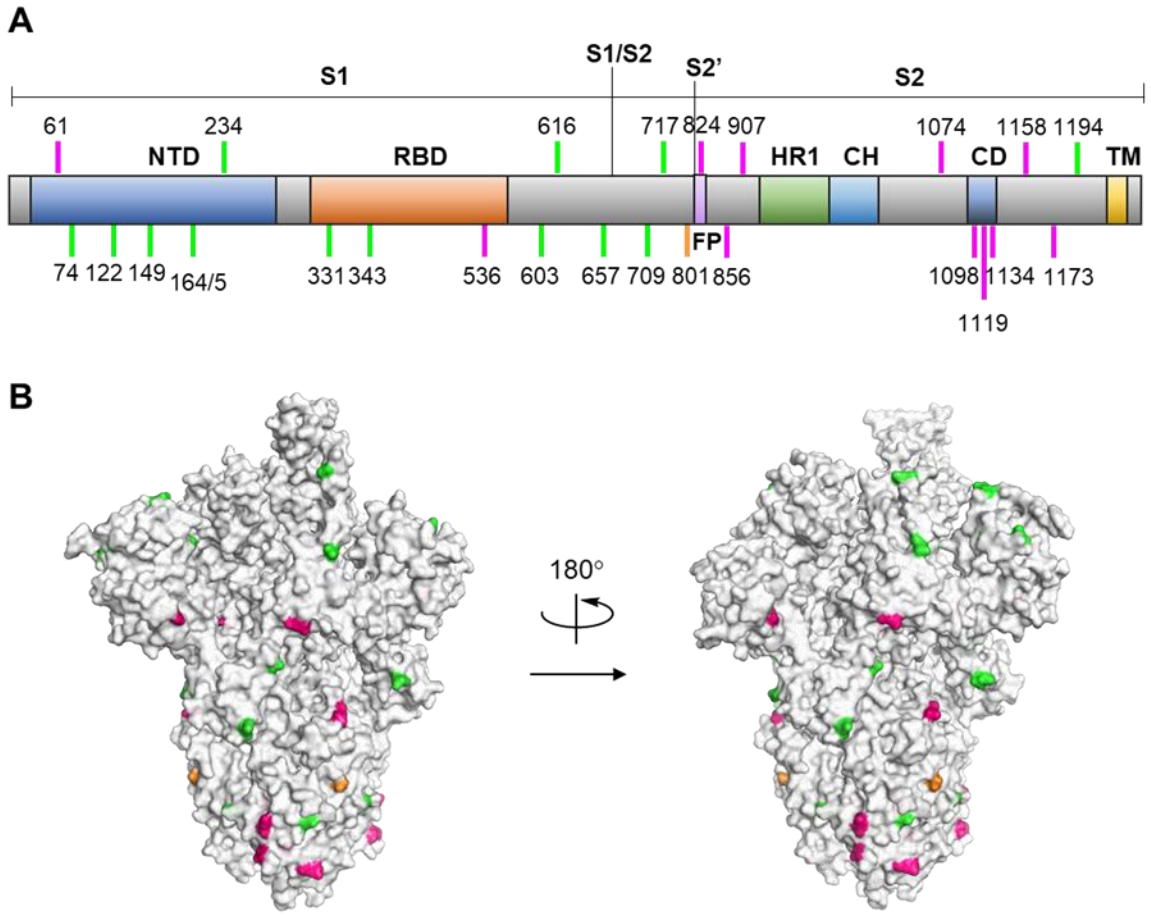


**Fig. S2. Glycosylation Pattern of the XBB.1.5 S(P2) Protein. A.** Schematic representation of the Omicron XBB.1.5 S glycoprotein. The positions of N-linked glycosylation observed by mass spectrometry studies are shown as bar lines. Glycan sites are colored according to glycan content: green (75 to 100%), orange (50 to 75%) and pink (0 to 50%). Protein domains are illustrated: N-terminal domain (NTD), receptor binding domain (RBD), fusion peptide (FP), heptad repeat 1 (HR1), central helix (CH), connector domain (CD), and transmembrane domain (TM). S protein S1 subunit, S2 (subunit after furin cleavage site to C-terminus) and S2' (subunit after TMPRSS2 cleavage site within the fusion peptide of Spike to C-terminus) boundaries are also illustrated. **B.** Structure-based mapping of N-linked glycans on the Omicron XBB.1.5 FL S(P2) Cryo-EM structure. Glycans are not modeled. Only glycosylated asparagine residues are highlighted and colored according to glycan content defined in **A**.

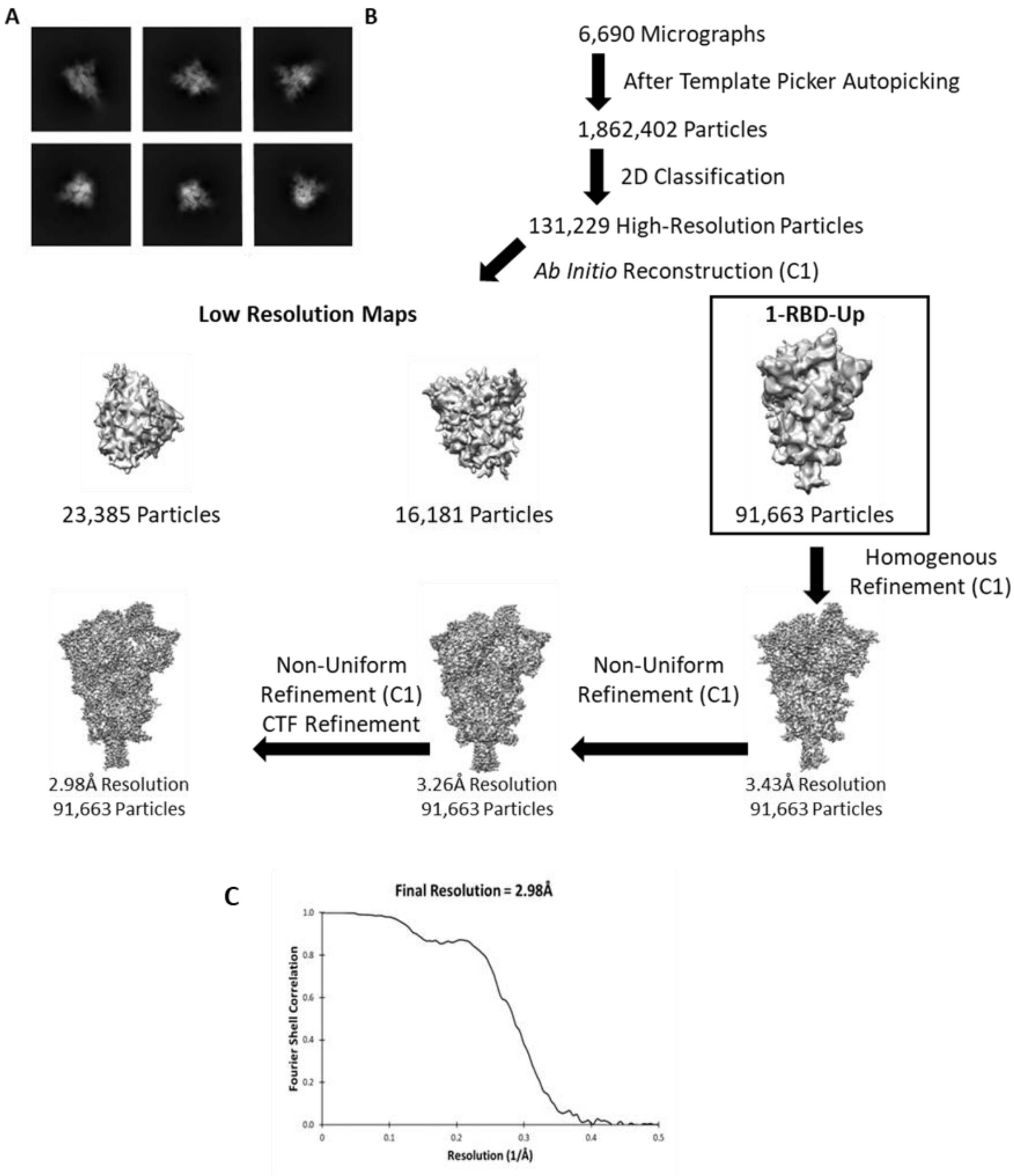


**Fig. S3. Cryo-EM workflow for construct pSB6534. A.** Representative two-dimensional class averages of Omicron XBB.1.5 FL S(P2). **B.** Cryo-EM data processing flowchart revealed 1-RBD-up as the primary S protein conformation. Particle number, Fourier shell correlation curve and corresponding resolution for the final map are indicated. **C.** Fourier shell correlation (FSC) curve indicates an overall nominal resolution of 2.98 Å using the gold standard FSC = 0.143 criterion.

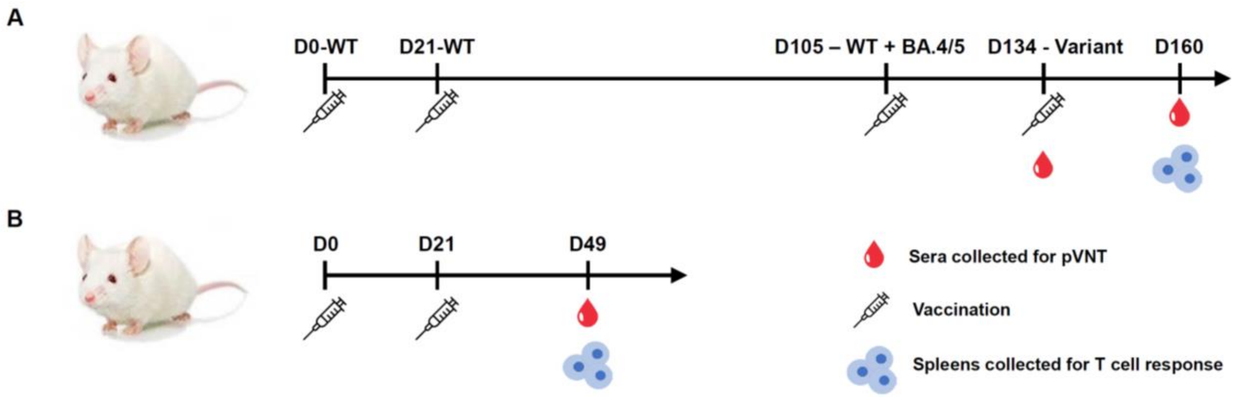


**Fig. S4. Schema for BNT162b2 variant-modified vaccine mouse immunogenicity studies. A.** In the booster study, female BALB/c mice were administered a two-dose series of the original monovalent BNT162b2 (WT) vaccine (D0, D21), followed by a third dose with the bivalent WT + BA.4/5 BNT162b2 vaccine (D105), and a 4th dose (D134) with one of the following variant-modified BNT162b2 vaccines:  monovalent Omicron BA.4/5, bivalent WT + Omicron BA.4/5, monovalent Omicron XBB.1.5 or bivalent Omicron XBB.1.5 + Omicron BA.4/5. Sera for pseudovirus neutralization test (pVNT) were collected just prior to the final vaccination (D134) and the last post-vaccination timepoint (D160). **B.** In the primary series study, female BALB/c mice (10 per group) were administered a two-dose series (D0, D21) of one of the following variant-modified BNT162b2 vaccines: monovalent Omicron BA.4/5, bivalent WT + Omicron BA.4/5, monovalent Omicron XBB.1.5 or bivalent Omicron XBB.1.5 + Omicron BA.4/5. Sera for the pVNT were collected at 2 weeks after the second vaccination (D49). In both **A** and **B**, mice were vaccinated in groups of 10, spleens were collected at the final post-vaccination timepoints, and the total dose level for each vaccine formulation administered was 0.5 µg.

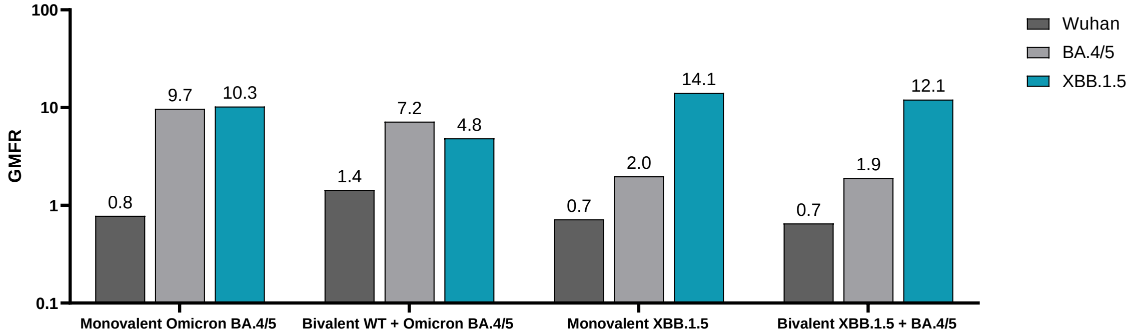


**Fig. S5. Geometric mean fold rise in pseudovirus neutralization titers ($NT_{50}$) from pre- to post-4th dose with a BNT162b2 Omicron XBB.1.5 variant-adapted vaccine in immune-experienced mice.** Female BALB/c mice (10/group) that were previously vaccinated with two-doses of original monovalent BNT162b2 (WT) vaccine , and one subsequent dose of bivalent WT + Omicron BA.4/5 vaccine received a single intramuscular booster dose of one of one of the following variant-modified BNT162b2 vaccines:  monovalent Omicron BA.4/5, bivalent WT + Omicron BA.4/5, monovalent Omicron XBB.1.5 or bivalent Omicron XBB.1.5 + Omicron BA.4/5. All vaccine formulations contained a total dose of 0.5 µg. Serum neutralizing antibody responses were measured by a pseudovirus neutralization assay. The fold rise in geometric mean neutralizing titers (GMFR) from pre-fourth dose to one-month post-dose are shown for the Wuhan reference strain, Omicron BA.4/5 and XBB.1.5. The limit of detection (LOD) is the lowest serum dilution, 1:20.

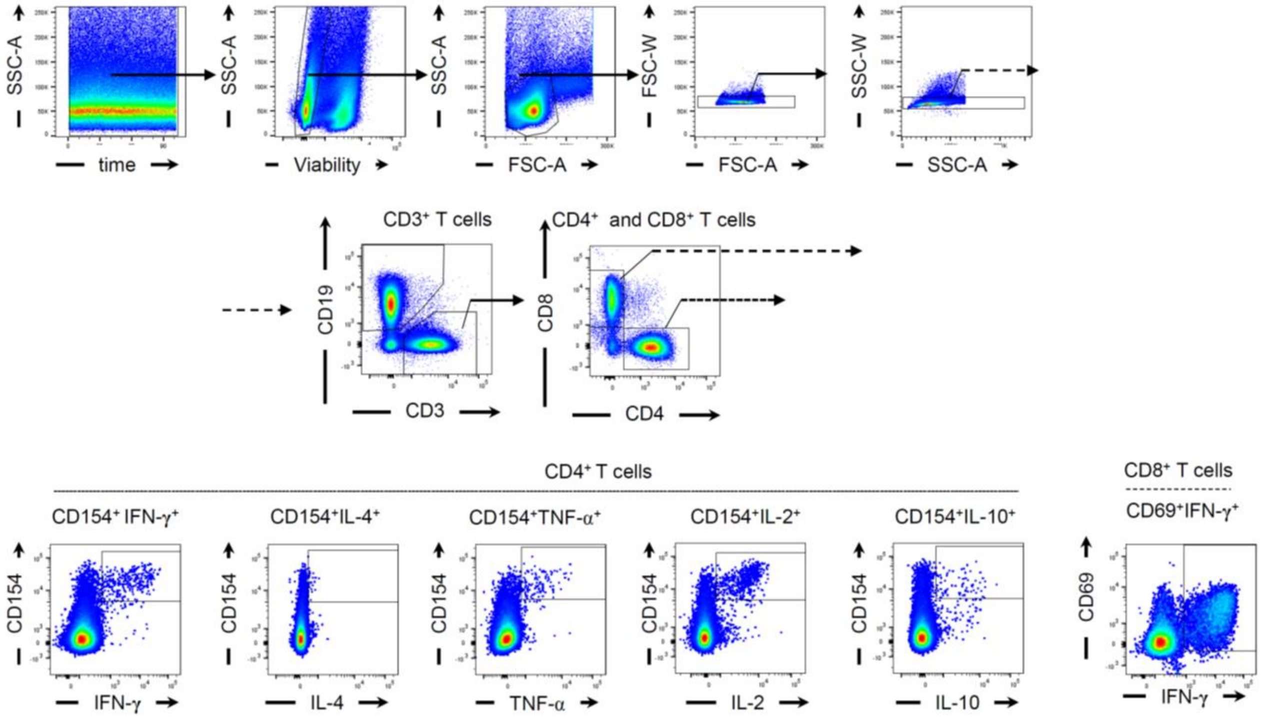


**Fig. S6. Gating strategy for intracellular cytokine staining flow cytometry analysis of T cell responses.** Flow cytometry gating strategy for identification of SARS-CoV-2 spike-specific T cells for different sublineages. (Upper row, left to right) Starting with events acquired with a constant flow stream and fluorescence intensity, viable cells, lymphocytes, and single events were identified and gated. Within singlet lymphocytes, CD19-negative CD3+ T cells were identified and gated into CD4+ and CD8+ T cells (middle row). Antigen-specific CD4+ T cells were identified by gating on CD154 and cytokine-positive cells. Activated CD8+ T cells were identified by gating on CD69 and cytokine-positive cells (bottom row). The antigen-specific cell frequencies were used for further analysis.

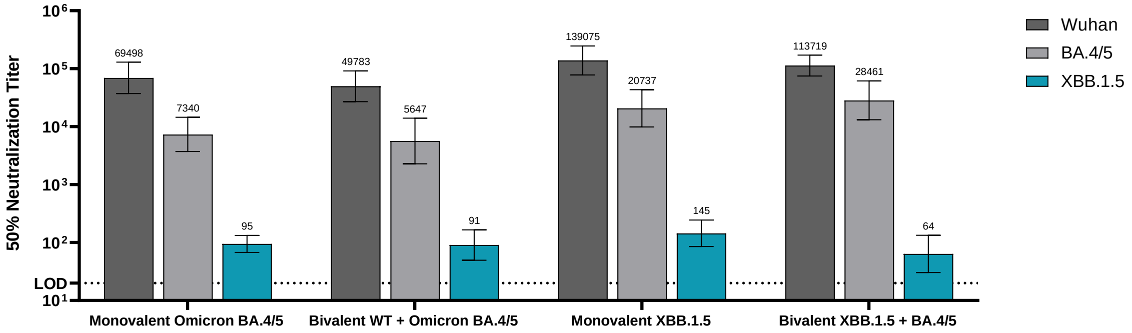


**Fig. S7. Baseline pseudovirus neutralization titers ($NT_{50}$) prior to BNT162b2 variant-adapted vaccine booster immunization in BNT162b2-experienced mice.** Female BALB/c mice (10/group) previously vaccinated with two doses of monovalent original BNT162b2, and one subsequent dose of bivalent BNT162b2 (WT + Omicron BA.4/5) received a single intramuscular dose of one of the following variant modified BNT162b2 vaccines: monovalent Omicron BA.4/5, bivalent WT + Omicron BA.4/5, monovalent Omicron XBB.1.5 or bivalent Omicron XBB.1.5 + Omicron BA.4/5. All vaccine formulations contained a total dose of 0.5 µg. Serum neutralizing antibody responses were assessed in a pseudovirus neutralization assay for the pre-fourth dose timepoint against the Wuhan reference strain, and the Omicron sublineages BA.4/5 and XBB.1.5. 50% pseudovirus neutralization titers are shown as geometric mean titers (GMT) ± 95% CI of 10 mice per vaccine group. The limit of detection (LOD) is the lowest serum dilution, 1:20.

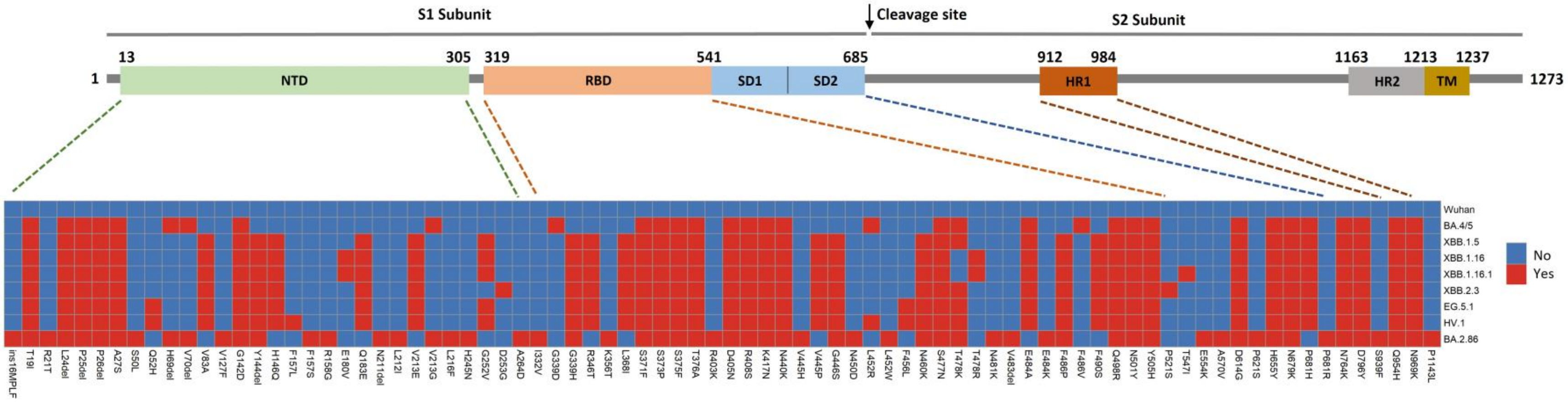


**Fig. S8. SARS-CoV-2 S amino acid sequence differences across lineages and sublineages for generated pseudoviruses.** Isolates metadata for each lineage was downloaded from GISAID (https://gisaid.org/) by filtering lineage name. Frequency of spike (S) protein amino acid difference relative to the ancestral strain within each lineage was calculated and a consensus S protein amino acid sequence list was generated by prioritizing sequence differences observed across more than 50% of sequences in the GISAID database. Consensus S protein amino acid sequence lists for all lineages were then aligned and plotted using R library heatmap. NTD – N-terminal domain; RBD-receptor binding domain; SD-subdomain 1; SD2-subdomain 2; HR1-heptad repeat 1; HR2-heptad repeat 2; TM-transmembrane domain.

**Table S1. Expression Constructs of FL S(P2) and RBD proteins**

| Code | Description | Mutations | Affinity Tag |
|---|---|---|---|
| pSB2782 | S(P2) (WT ) | K986P, V987P | C-terminal TwinStrep |
| pSB6534 | S(P2) (Omicron XBB.1.5) | K986P, V987P, T19I, L24del, P25del, P26del, A27S, V83A, G142D, Y144del, H146Q, Q183E, V213E, G252V, G339H, R346T, L368I, S371F, S373P, S375F, T376A, D405N, R408S, K417N, N440K, V445P, G446S, N460K, S477N, T478K, E484A, F486P, F490S, Q498R, N501Y, Y505H, D614G, H655Y, N679K, P681H, N764K, D796Y, Q954H, N969K | C-terminal TwinStrep |
| pSB2661 | RBD (WT ) | | C-terminal StrepII |
| pSB6876 | RBD (Omicron XBB.1.5) | G339H, R346T, L368I, S371F, S373P, S375F, T376A, D405N, R408S, K417N, N440K, V445P, G446S, N460K, S477N, T478K, E484A, F486P, F490S, Q498R, N501Y, Y505H | C-terminal His×8 |

**Table S2. Cryo-EM Data Collection, Processing and Refinement Statistics**

| **Data Collection and Processing** | Omicron XBB.1.5 FL S(P2) |
|---|---|
| Magnification | 130,000x |
| Voltage (kV) | 300 |
| Total Electron Exposure ($e^-/Å^2$) | 40.0 |
| Defocus Range (μm) | -0.6 to -2.8 |
| Pixel Size (Å) - Counting Mode | 0.750 |
| Movies Recorded | 6,690 |
| Symmetry Imposed | C1 |
| Initial Particle Images (no) | 1,862,402 |
| Final Particle Images (no) | 91,633 |
| Map Resolution at FSC=0.143 (Å) | 2.98 |
| **Refinement** | |
| Map sharpening *B* factor ($Å^2$) | -55.8 |
| Model composition in the asymmetric unit | |
| Non-hydrogen atoms | 21,179 |
| Protein residues | 2,758 |
| B factors ($Å^2$) | |
| Protein | 35.997 |
| R.M.S. Deviations | |
| Bond lengths (Å) | 0.004 |
| Bond angles (°) | 0.734 |
| Validation | |
| MolProbity score | 1.78 |
| Clashscore | 3.04 |
| Poor rotamer (%) | 2.71 |
| Ramachandran Plot | |
| Favored (%) | 94.83 |
| Allowed (%) | 5.17 |
| Outliers (%) | 0.00 |
